# Dynamic membership drives long-term persistence of sparrow winter social communities

**DOI:** 10.1101/2025.07.25.666472

**Authors:** Anastasia E. Madsen, Bruce E. Lyon, Alexis S. Chaine, Daizaburo Shizuka

## Abstract

Social groups range from highly stable to highly dynamic over extended periods. Demographic factors like individual lifespans and rates of births, deaths, immigration, and emigration in a population can strongly impact group stability and persistence over time. Highly stable long-term groups are often driven by long-lived core individuals that maintain relationships, kin-structure, or high philopatry. In opposition, fluid groups of individuals that associate repeatedly, called social communities, can persist in populations that have high turnover, lack kin structure, and exhibit high fission-fusion dynamics. However, we lack an understanding of how animal social communities persist amidst highly dynamic membership. We investigated the drivers of persistence of social communities in a winter population of migratory golden-crowned sparrows over 12 years. Sparrows frequently flock with the same individuals over the entire winter, forming social communities that are identified from discrete clusters within a social network. Long-term community membership was highly dynamic due to high annual turnover and the relatively short lifespans of individuals. However, the social fidelity of returning members and recruitment of new members between consecutive years resulted in communities that persisted for multiple years, even when there was a complete turnover of members over time. Our results indicate that groups can persist with consistent short-term membership, overlapping generations, and turnover with balanced rates of recruitment and loss. This dynamic stability of individual membership may be a universal feature of large, long-term groups found in many social systems, from animal societies with fission-fusion dynamics to networks of human collaboration.

## Introduction

Animal populations are often organized into social groups in which individuals maintain differentiated patterns of relationships. Individuals can gain fitness benefits from remaining in the same group, but may leave when social or ecological conditions change.^1,2^ In nature, groups range from relatively small, stable, and cohesive in space and time to large and dynamic.^2,3^ Importantly, stability and dynamism in group membership can occur at different temporal scales, which is well-exemplified by groups with fission-fusion dynamics.^4–8^ Such groups have smaller sub-groups with dynamic short-term membership, yet maintain stable relationships over longer periods, forming distinct “social communities” consisting of individuals that associate more frequently with each other than with others in the population.^3,9–12^ Crucially, demographic turnover—i.e., births, deaths, immigration, and emigration—directly and permanently alters group membership.^13^ Social groups may destabilize after key members are lost,^14–16^ succumb to demographic stochasticity after losing many members^17^ (an especially important consideration for small groups), or reorganize after gaining new members.^18–20^ Social groups play a central role in the ecology and evolution of many animals,^21–26^ yet we rarely grapple with a fundamental puzzle for their long-term existence: can social groups persist beyond the lifetime of their members, and if so, what determines the longevity of a social group when members inevitably turnover through deaths, births, and immigration?

Different dimensions of life history can drive stability and persistence of social groups, such as lifespan, overlapping generations, and philopatry.^13,27^ Slow life histories may foster stable groups; for example, the long individual lifespans of African savanna elephants and Southern orcas result in long-lasting relationships, and subsequently, long-lasting groups.^28–30^ Social groups can span multiple generations due to the roles of elder members (e.g., matriarchs), including the maintenance of long-term relationships within and between groups, knowledge transmission, and inheritance of social positions.^16,31–35^ Moreover, philopatry can help maintain social group stability by anchoring individuals in familiar social and ecological contexts.^36–39^ Mechanisms like high genetic relatedness, extended parental care, delayed dispersal, and sex-biased philopatry in kin groups can support stability and persistence based around consistent core members.^40–48^ These stable components of life history can facilitate consistent interactions and enduring group cohesion, leading to high stability and persistence of social groups.

Research on social group stability and life history has largely focused on long-lived animals with cohesive social groups characterized by high philopatry and kin structure. However, a major gap remains in our understanding of the persistence of less cohesive social groups that are common in animals with short lifespans, no kin structure, or fission-fusion dynamics that result in a high turnover of members. The application of social network analysis across such species (e.g., small rodents,^49–51^ sharks,^52,53^ and passerine birds^54–57^) has revealed that fluid social associations are often clustered into social communities of individuals that associate frequently with each other. Importantly, these social communities can persist despite high annual turnover of membership^54^ or large mortality events^49^ and can outlast the lifespans of their members.^49^ These observations again point to the fundamental question: under what conditions can groups be stable while membership is highly dynamic? One clue to resolving this puzzle comes from studies of group dynamics in human networks,^58–60^ which show that the rate of membership turnover can influence the long-term persistence of groups. Dynamic membership may disrupt internal group dynamics,^59,61^ but as individuals inevitably leave, new recruits are necessary to sustain groups in the long term^58,59,62^. Thus, for groups to persist over long timespans, they must be able to tolerate some level of turnover. Here, we use a 12-year dataset on winter flock networks of a migrating songbird, the golden-crowned sparrow (*Zonotrichia atricapilla*), to ask how the turnover of membership through demographic processes such as deaths and immigration lead to the persistence or loss of social communities in a natural population.

Our long-term study on golden-crowned sparrows shows that these birds maintain seasonal social relationships at their wintering grounds despite their short lifespans and long-distance migrations.^54^ On the wintering grounds, golden-crowned sparrows forage in flocks that show high fission-fusion dynamics. Flocks move cohesively across the landscape for short periods of time, but individuals join and leave flocks within and between days. At the population scale, repeated associations form social networks composed of distinct communities of birds that flock together often. These winter social relationships are not based on kinship^63^ and are exclusively seasonal—all sparrows migrate, and there is no evidence that winter relationships are maintained in migration or on breeding grounds.^64^ Furthermore, the turnover of individuals at the wintering site between years is high, with typically half of the birds not returning between years.^65–67^ Nevertheless, pairwise social associations between returning birds across years are consistent, indicating that birds continuously form flocks with the same flockmates (i.e., birds that frequently flock together, as opposed to mating pairs).^54,65^ These stable pairwise interactions also exert a strong influence on space use. Although many returning birds have high site fidelity (average home range shift = 27.3m), this fidelity can be weakened by the loss of many close flockmates, signaling a strong effect of social fidelity on fine-scale space use.^68^

Previously, we found that many surviving sparrows returned to the same social communities across the first three years of this study.^54^ However, membership in social communities does not remain entirely static. The composition of communities can change as individuals join the population, leave the population, or move to a new community. These changes to sparrow community composition lead to variation in structural properties like cohesion (i.e., proportion of social connections within vs. outside the community).^65^ With dynamic community structure and composition and high turnover rates, we might presuppose that stable sparrow communities are short-lived. However, this variability in structure and composition also provides an ideal opportunity to investigate the social and demographic factors that underlie temporal stability of social communities. This system thus allows us to ask: why do some social communities disappear or split, while others persist? To tackle this question, we apply a social network approach developed to ask similar questions about the temporal dynamics of community membership in other systems (e.g., human collaboration).^58,69^ In this approach, we track the individual membership in communities and quantify the changes in composition over time. This allows us to investigate the causes and consequences of *stationarity* and *persistence* of social communities. Here, stationarity refers to the consistency of community membership across time, and persistence refers to the duration of time over which a social community maintains overlapping core membership (as opposed to a community losing all of its members or members splitting into separate communities between time points). We show that golden-crowned sparrow communities had highly nonstationary (i.e., dynamic) *long-term* membership, but the joint overlap of members between consecutive years and the gradual recruitment of new members led to persistent social communities that outlived all of their original members.

## Results

### Community Detection

Using flock observations, we quantified social relationships over 12 winters using the simple ratio index (SRI), which reflects the pairwise strength of association between birds.^70–73^ These associations are the edges, or connections, in each social network. Unlike our previous work, we included first-year immigrants in the networks to understand the effects of recruitment on social communities. For each of the 12 winter networks, we used the Louvain community detection algorithm to identify clusters of individuals with more connections within a community than between communities.^12^ Using this definition, there was an average of 3.9 (SE±0.23) social communities in each year in the study population.

To determine whether communities persisted over time, we used a dynamic community detection algorithm, *MajorTrack*, to track community membership between each consecutive annual network.^69^ This method determines whether a community is the same between time points by considering only the individuals that are present in both networks being compared. A social community that retained a majority of the same returning members between consecutive networks was considered the same (see methods for additional details).^69^ We detected 17 unique communities across the study period (Figure 1). On average, communities persisted for 2.6 (SE±0.62) years, while the average individual sparrow lifespan was 1.6 (SE±0.04) years. The longest-lived community spanned 8 years, outlasting all original members.

**Figure 1.**
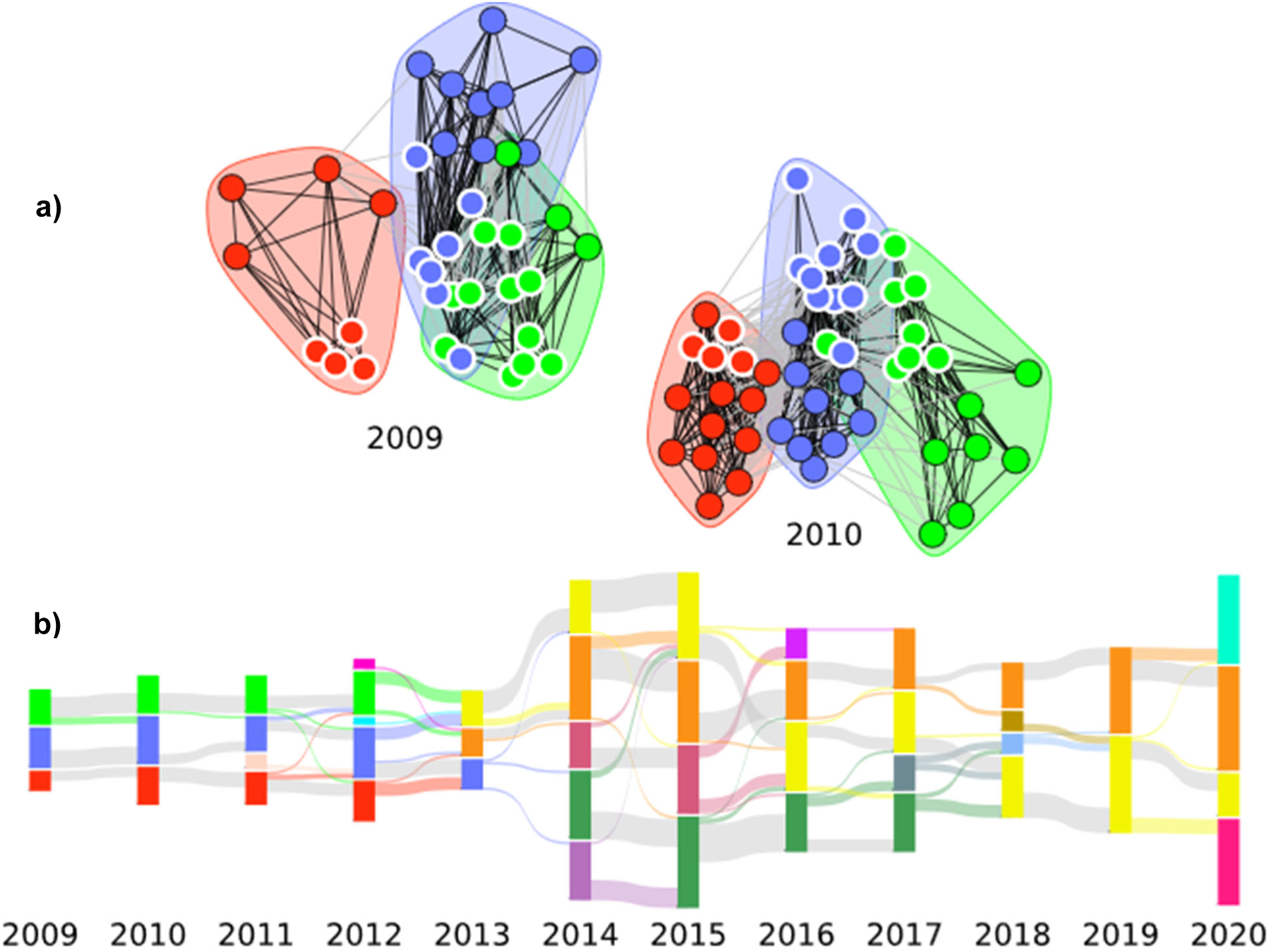
Dynamic community membership of golden-crowned sparrows from 2009-2020. Colors correspond to a unique social community identity that was detected across consecutive networks. **a)** Networks from 2009 and 2010, with clusters and nodes colored according to their identity. Nodes that are present in both networks are outlined in white. **b)** Alluvial plot showing the flow of membership through communities over time. Vertical stacks represent social communities within a year, and horizontal bands represent the movement of individuals to social communities between years. The color of horizontal bands corresponds to the source community, while gray horizontal bands represent individuals returning to the same community between years. The height of stacks is relative to the size of communities. The height of the vertical stacks and horizontal bands differ when individuals have entered the population for the first time (no left flow) or left the population (no right flow). *N* = 567 total birds and 17 communities.

### Community Stationarity

We assessed the consistency of social community membership (i.e., stationarity) by first correlating community membership between years. The membership correlation is the ratio of the consistent members to all unique members in a community between two years (see methods, Eqn. 1).^58^ The correlation includes the lost members and new recruits in the denominator, allowing us to account for changing group size and to represent the full dynamism of membership over time. To evaluate the change in correlations over the lifespan of a community, we included only communities that persisted for at least three years in our analysis (yielding a minimum of two correlation values for a community), with a final sample of six multiyear communities (*N*=27 correlation values).

We first calculated the correlation in community membership between each consecutive pair of years that a community persisted (*t*, *t_+1_*), which we call *C*. *C* represents how similar membership is between years amidst turnover by capturing the retention, loss, and gain of members from one year to the next. Large values indicate low turnover, i.e., few losses and low recruitment, while small values indicate many losses, high recruitment, or a combination of both. We might expect that long-term communities will retain many core members from the previous year, corresponding to high *C* values over time, or that membership turnover rates are stable over time, corresponding to constant *C* values over time. However, this correlation was low for many communities (mean *C* = 0.2), indicating low stationarity. *C* surprisingly decreased over the course of community lifespans (Figure 2a, Table 1).

**Figure 2.**
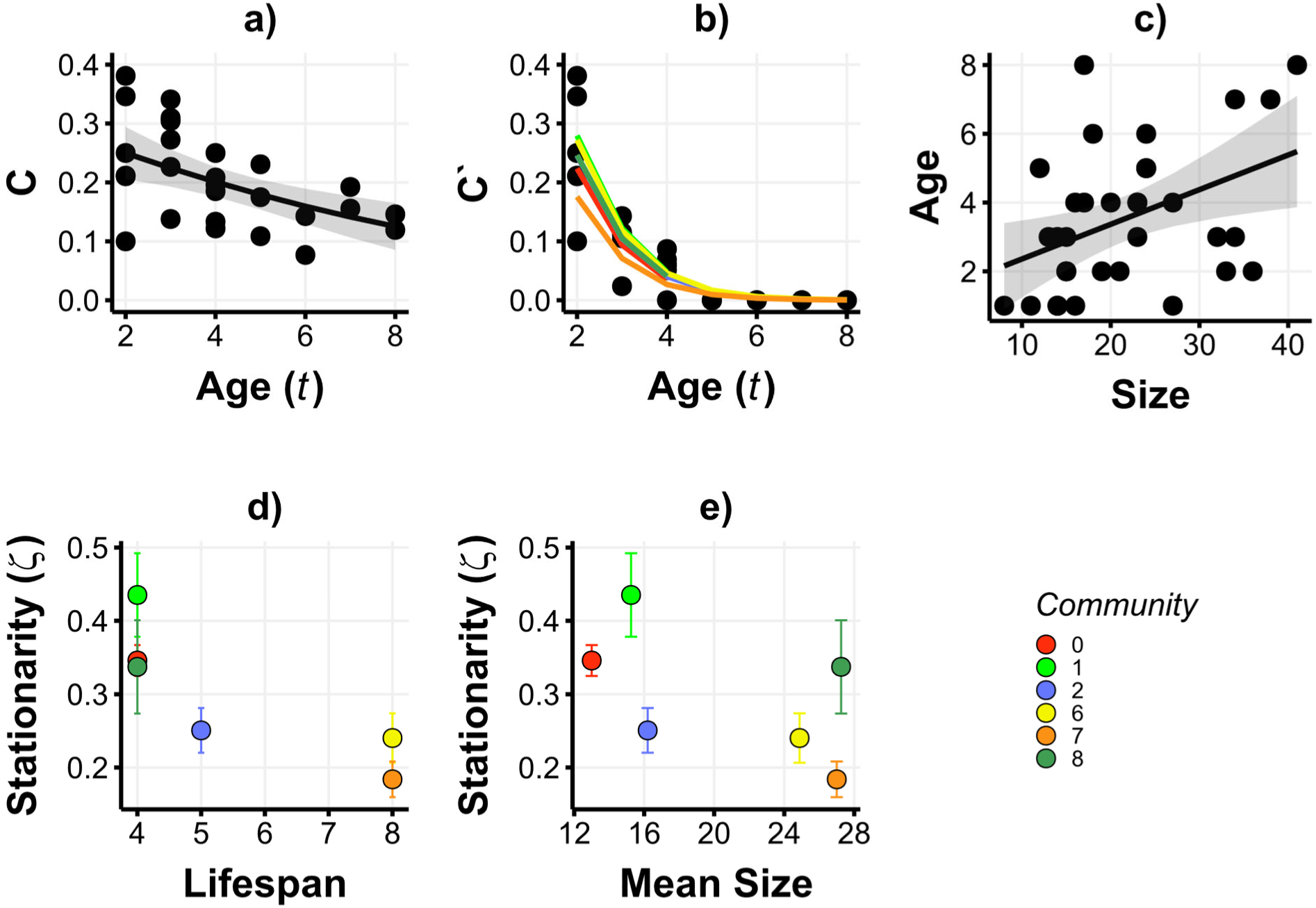
Dynamic membership of long-term communities. For each social community that persisted for at least 2 years (*N* = 6 communities, each shown in a different color), we show across the top row **a)** the correlation in membership (*C*) between consecutive years (*t*, *t+1*), **b)** the stability in membership (*C’*) between subsequent years (*t_0_*, *t_0_*+*t*) as a function of community age (*t*), and **c)** community age as a function of size. Bands around the regression line show the standard errors of regression models. The bottom row shows **d)** the index of stationarity (𝜻) for communities as a function of community lifespan (*t_max_*) and **e)** 𝜻 as a function of mean community size. Error bars represent the standard error for 𝜻.

**Table 1.** Long-term and consecutive-year community memberships are dynamic. Binomial regression models were used to estimate the relationship between the age of a community and the correlation of consecutive year membership, *C*, and the age of a community and the correlation of original membership, *C’*. The minimum community age in these models was 2 because both *C* and *C’* represent pairwise comparisons between years. *C* was found to linearly decrease with age and *C’* exponentially decreased with age (*N*=27).

| Model | Variable | Estimate | SE | P-value |
| --- | --- | --- | --- | --- |
| $C \sim \text{Community Age}$ | Intercept | -0.82 | 0.19 | <0.001* |
|  | Community Age | -0.14 | 0.043 | 0.001* |
| $C' \sim \text{Community Age}$ | Intercept | 0.88 | 0.43 | 0.041* |
|  | Community Age | -1.02 | 0.14 | <0.001* |

**Table 2.** Community size is correlated with its age. A linear regression model showed a significant positive correlation between the size and age of a community (*N*=33).

| Model | Variable | Estimate | SE | P-value |
| --- | --- | --- | --- | --- |
| Community Age ∼ Size | Intercept | 1.35 | 0.92 | 0.15 |
|  | Size | 0.10 | 0.039 | 0.016* |

Next, we calculated the correlation in membership between the first year (*t_0_*) and each subsequent year (*t*) that a community persisted, which we call *C’*. Similar to *C*, this evaluates the similarity in membership while explicitly investigating membership turnover across time. *C’* represents the turnover of membership from the birth of a community to each subsequent year until its disappearance (Eqn. 2). We found that *C’* decreased exponentially as the age of a community increased, as may be expected because the population turnover rate is close to 0.5^68^ and *C’* is limited by short individual lifespans (mean = 1.6 years; Figure 2b). In a striking result, some communities persisted beyond the lifespan of all original members, eventually experiencing complete membership turnover (Figure 2b, Table 1). Additionally, we evaluated the influence of group size on community age, considering that small communities may be more vulnerable to demographic stochasticity. Importantly, community age represents how many years that a community has existed, not the age of individual members in a community. As expected, we found that larger communities were significantly older than smaller communities (*N*=33, Figure 2c, Table 1).

Finally, to evaluate the overall stationarity of each community over the entire timespan they persisted, we calculated a stationarity index (𝜻) for each community (Eqn. 3, see methods).^58^ The stationarity index gives a single value for each community, representing the average correlation in consecutive community membership over the entire community lifespan, with higher values representing communities with more stable membership and lower values representing communities with more dynamic membership. The total membership stationarity, 𝜻, was relatively low across communities, averaging 0.29 (SE±0.037). However, counter to expectations, communities with the lowest stationarity indices generally persisted longer than those with higher stationarity (Figure 2d). Additionally, the largest communities were on average the least stationary, indicating that large communities can withstand highly dynamic membership (Figure 2e).

## Discussion

We found that golden-crowned sparrow populations can have social communities that persist over multiple years, outliving any single sparrow in a community. Social systems with high fission-fusion dynamics are inherently dynamic at the smallest time scales, but the patterns of repeated associations between individuals can result in stable structures over longer periods of time. Despite that individual sparrows only associate for a few months of the annual cycle, that the social context lacks many established drivers of social stability,^63,64^ and that the population experiences high turnover rates,^66^ we found surprisingly long-lasting social communities.^54^ By analyzing the persistence of dynamic communities in a natural population, our study provides an approach for studying the causes and consequences of long-term stability of groups in a broader array of study systems that lack classic hallmarks of social groups such as kinship and cooperation.

The persistence of social communities relies on two key demographic processes: retention of old members and recruitment of new members that keep pace with the loss of members. Because golden-crowned sparrows are limited by short lifespans (mean = 1.6 years), retaining members cannot solely account for the long-term persistence of social communities. Indeed, the longest persisting community in our population spanned 8 years, exceeding the lifespan of the average sparrow and the membership of any single individual (Figure 2b). Some degree of overlapping membership is necessary for communities to persist, and many returning birds rejoin communities with their flockmates from the previous year (gray horizontal bands, Figure 1b). However, this overlap was surprisingly low relative to the total number of members that were lost and recruited between years (Figure 1, Figure 2a). Contrary to expectations, high membership stationarity is not necessary for the long-term persistence of communities, and in fact, the consecutive membership correlation, *C*, significantly decreased over time (Figure 2a, Table 1). Furthermore, community size influenced its longevity: large communities generally persisted longer than small communities (Figure 2c, 2d). This is consistent with the expectation that smaller communities are more prone to disappearance due to demographic stochasticity.^40^ Small communities also tended to have higher overall stationarity, meaning their membership was less dynamic than large communities (Figure 2e). This pattern suggests that large communities may be more successful at recruiting new members or else better able to buffer the effects of high turnover.^40,58^ These results align with properties of other social communities, including parallels in humans. Many human social communities experience high turnover rates,^58,59^ with communities like academic societies and institutions frequently surviving the complete turnover of members.^58^ Across human and animal systems, social communities persist when they can successfully recruit new members and balance turnover rates.^58,59^ This suggests that dynamic membership due to turnover, particularly recruitment, may be a general mechanism that allows long-term social groups to persist in nature.^40,74^

Our study highlights a fundamental question about the persistence of social groups in nature: how and when do social groups persist beyond the lifespan of individual members? Turnover of membership is an inherent process underlying all social groups across time, as it is with social systems more generally.^13,74–77^ Yet, each individual loss or gain of a member does not immediately lead to the breakdown of social groups. In some species, social groups can persist beyond all individual lifespans because individuals can socially inherit a breeding territory, breeding position (e.g., cooperative breeding groups),^39–41^ social connections,^31,78^ or dominance status.^32^ However, we find that long-term persistence can also occur in animal social groups with high turnover and that lack such processes of social inheritance. This suggests that the impact of membership turnover on social groups varies based on the consequences of turnover on group dynamics.^79–82^ In some cases, high rates of turnover could be destabilizing,^83^ for example, in systems with highly cohesive groups that rely on coordination and cooperation to persist.^34,44,80,84,85^ Meanwhile, groups with high fission-fusion dynamics like social communities may persist because they can respond flexibly to turnover and more quickly replenish members with new recruits.^20,58^ Depending on the system, intragroup dynamics could determine the magnitude of turnover that a group can withstand, or contrarily, the demographic turnover rate could be a fundamental driver of social group dynamics. ^74,76,77,86^ Moreover, feedback between group dynamics and turnover, life history, and socioecological dynamics all contribute to the persistence of social groups in complex ways that necessitate system-specific considerations.^87–89^

We provide evidence that the rate of demographic turnover is a key driver of long-term social dynamics, and critically, that dynamic membership can permit the long-term persistence of fluid groups like social communities. Communities can be resilient to demographic turnover because individuals are able to change and adapt to the dynamic social environment. The persistence of communities prevail as an unexpected, or emergent, outcome of consistent individual interactions.^87,88,90–93^ This is characteristic of a resilient complex adaptive system, which has structures that can both maintain stability and allow change, relative to their respective scales.^4,8,94,95^ Understanding the limits of the dynamic stability of social systems with high turnover has implications for the evolution of social dynamics that lead to stable social communities,^96^ facilitate cultural evolution, ^88,97–99^ and promote the development of complex social signals.^100–103^

## Resource Availability

Requests for further information should be directed to the lead contact, Anastasia Madsen. This study did not generate new unique reagents. All data and original code have been deposited at Data Dryad (DOI: 10.5061/dryad.bcc2fqznh) and will be made publicly available as of the date of publication.

## Acknowledgements

This work was supported by the University of Nebraska, the University of California, and the National Science Foundation (NSF) IOS-1750606. A.E.M. was supported by NSF DGE-1735362 and NSF DBI-2305860. A.S.C. was supported by the Laboratoire d’Excellence (LABEX) TULIP (ANR-10-LABX-41). B.E.L. was supported by a University of California, Santa Cruz Special Research Grant. We thank the UC Santa Cruz Arboretum staff for the use of facilities and thank the multitude of staff, volunteers, and students who assisted in collecting flock data throughout the duration of the study.

## Author Contributions

Conceptualization, A.E.M. and D.S.; Formal analysis, A.E.M.; Writing – original draft, A.E.M.; Writing – review and editing, A.E.M., D.S., A.S.C., and B.E.L.; Investigation, D.S., A.S.C., and B.E.L.; Data curation, D.S.; Funding acquisition, A.E.M., D.S., and B.E.L.

## Methods

### Data collection

Golden-crowned sparrows were banded at the University of California Santa Cruz Arboretum from 2009-2020 with unique combinations of color bands for individual recognition. Birds were caught in potter traps baited with millet seed and released after banding at the capture site.

Each winter, we collected flock observations beginning in October when most birds had arrived at the winter site and ending in March when birds began to break up winter flocks and start migration to breeding grounds. We determined flock membership by identifying all individuals within a 5m radius of each other and recording their presence on a 10m^2^ grid.^54^ Flocks were observed until all individuals were identified. Observations of different flocks were considered independent if observations were separated by at least 20 minutes.

#### Community detection

Memberships in foraging flocks were used to build a group-by-individual matrix for social network analysis for each of the 12 winter seasons. We calculated association matrices for each winter using the simple ratio index (SRI), which represents the proportion of observations in which two individuals were both present in a flock out of all recorded observations, both observed separately and together (R package *asnipe*) ^70–73^.

Importantly, membership in a flock does not automatically designate membership in a community, as community membership represents a summary of all flocks in a season. We used a hierarchical clustering algorithm (Louvain clustering, R package *igraph*),^104^ to identify community clusters for each winter network. Communities identified in this way represent a collection of individuals that interact more with other individuals within a community than with individuals in other communities. Using these community assignments, we used a post-hoc algorithm, MajorTrack, that detects dynamic communities by evaluating similarity in membership of communities between two networks.^69^ The algorithm calculates the proportion of community members that are present in consecutive network clusters, considering only the members that returned to the winter site, and therefore were present in both sets (e.g., the highlighted nodes in Figure 1a). Importantly, this allowed us to identify communities based on membership while also flexibly accounting for the gain and loss of individuals from the population between years. Communities in consecutive network clusters that contained a majority of the same members were considered the same multiyear community (Figure 1). A new social community was detected when it was composed entirely of new immigrants, when multiple communities merged into one, or when a community split into two or more communities. A community retained its identity when it gained only a small minority of members from a different pre-existing community; in this case, a merge event resulted in the loss of a community. The resulting community modules are visualized using an alluvial plot, which shows the dynamic membership of social communities between consecutive years (Figure 1b).

#### Community stability

To assess the stability of communities over time (*C*), we compared community membership, *A*, between each consecutive year the community persisted (*t*, *t+1*):

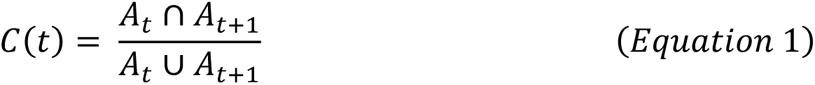

We evaluated how *C* changed over the lifespan of communities using a generalized linear mixed effects model with a binomial logit link function. We modeled *C* as a binomial response, which allowed us to explicitly account for differences in community size by modeling the raw counts of the same members and different members. Community age (*t*) was the predictor variable and the community identity was fit using a random slope and intercept (R package *lme4*)^105^. However, this model showed a variance of zero between the random effects and a singular model fit, indicating the model was likely overfit with too many random effects being specified. An additional model with only random intercept specified also resulted in a singular model fit, so no random effects were specified in the final model.

Next, we compared community membership, *A*, between the “birth”, or first year a community was seen (*t_0_*), and each subsequent winter it persisted (*t*).

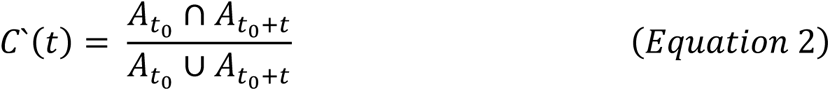

We evaluated how *C’* changed over the lifespan of communities using a generalized linear mixed effects model with a binomial logit link function, specifying *C’* as a binomial response, as with *C* above. Community age (*t*) was the predictor variable and community identity was fit with a random slope and intercept (R package *lme4*)^105^. This model was also overfitted, so community identity was only specified with a random intercept in the final model.

We then calculated an index of stationarity (𝜁) for consecutive year membership correlations, *C*, over the lifespan of a social community (*t_0_*:*t_max_*) to assess the stability versus dynamism of membership for each community.^58^

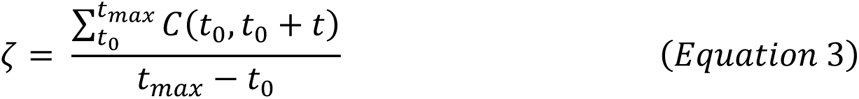

The stationarity index, 𝜁, represents how similar interannual membership is over the lifespan of each social community, where 0 represents complete turnover of members, and 1 represents completely static membership over time.

Finally, we investigated whether community size influences community age using a generalized linear mixed model with community identity as a random intercept. This model indicated a singular fit, and year was then fit as a random intercept instead; however, this model had significant quantile deviations, suggesting that the model was overfit. The final model we selected is a simple linear regression with community age as the response and community size as the predictor.

All model assumptions were checked by assessing the residuals of linear and generalized linear mixed models in the R package DHARMa.^106^ The final models met linear model assumptions with no significant quantile deviations, no overdispersion, and uniform simulated residuals (see Figures S1-S4).

