## Supplemental Information for "Dynamic membership drives long-term persistence of sparrow winter social communities"

Document S1. Figures S1-S3

This supplement contains the results of model assessment for the final models that were reported in the main text. We used the R package DHARMa to assess model residuals. We evaluated each model for significant quantile deviations, overdispersion, and uniformity of residuals. We used the simulateResiduals function in DHARMa to calculate scaled residuals from each model and to test for significant quantile deviations from simulated data. There were no significant deviations (Figures S1-S3).

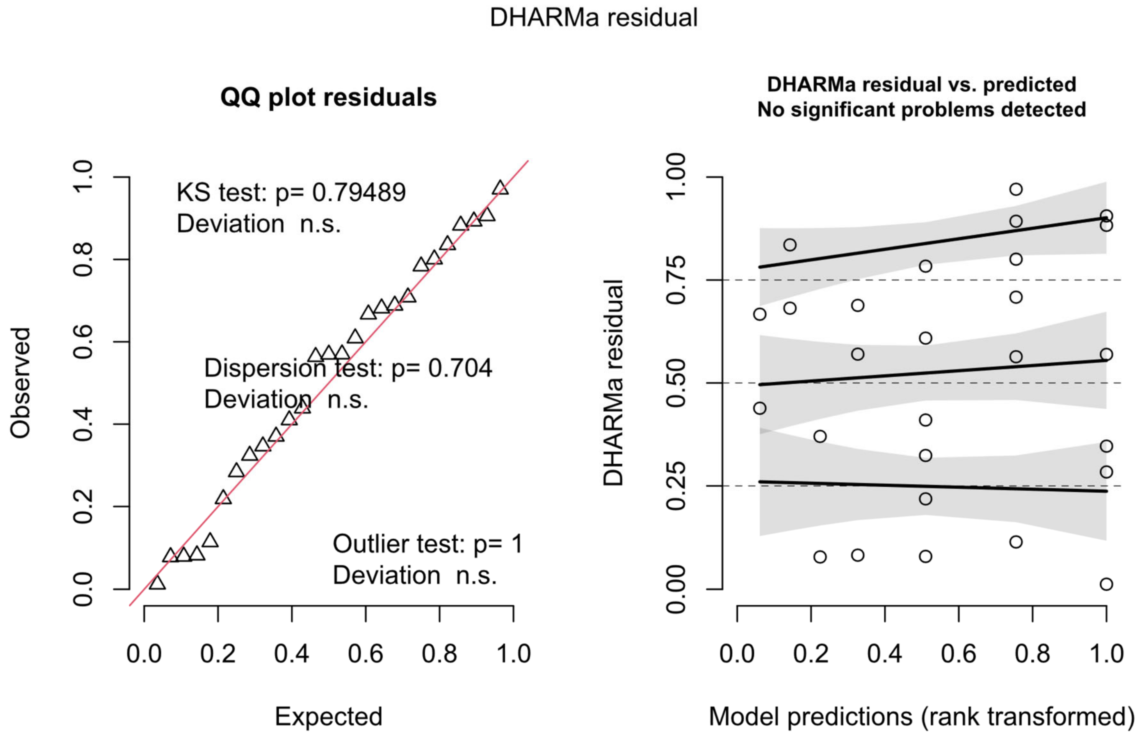


Figure S1. DHARMa simulated residuals for the final consecutive membership correlation (*C*) model, related to Table 1.

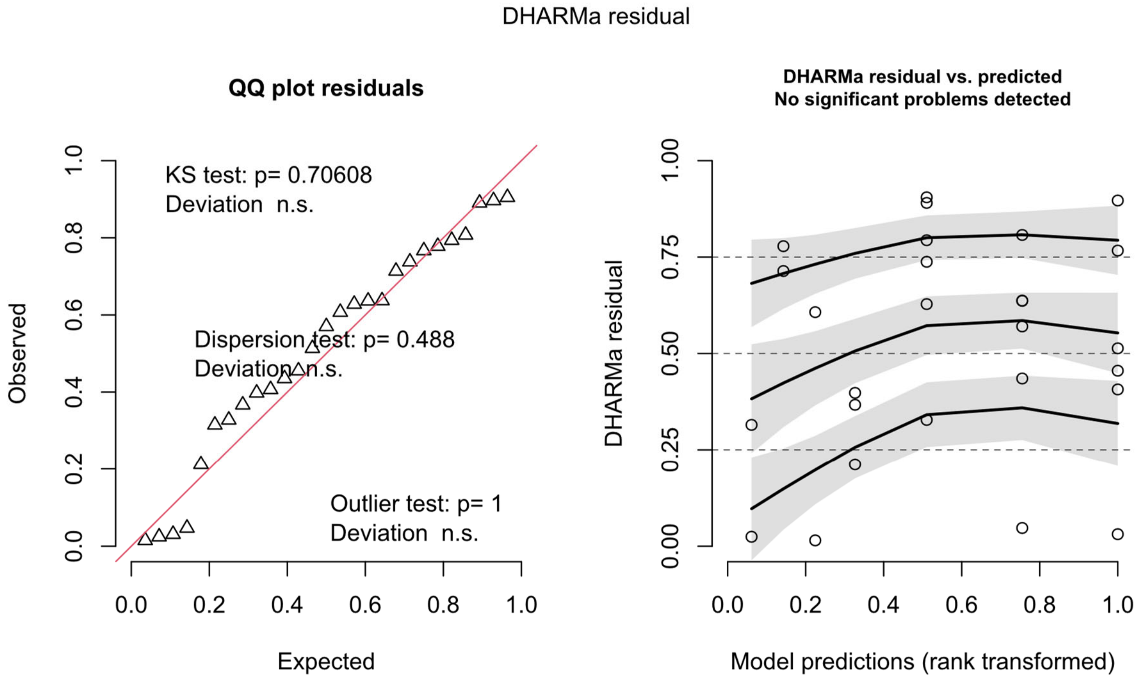


Figure S2. DHARMa simulated residuals for the final subsequent membership correlation ($C`$) model, related to Table 1.

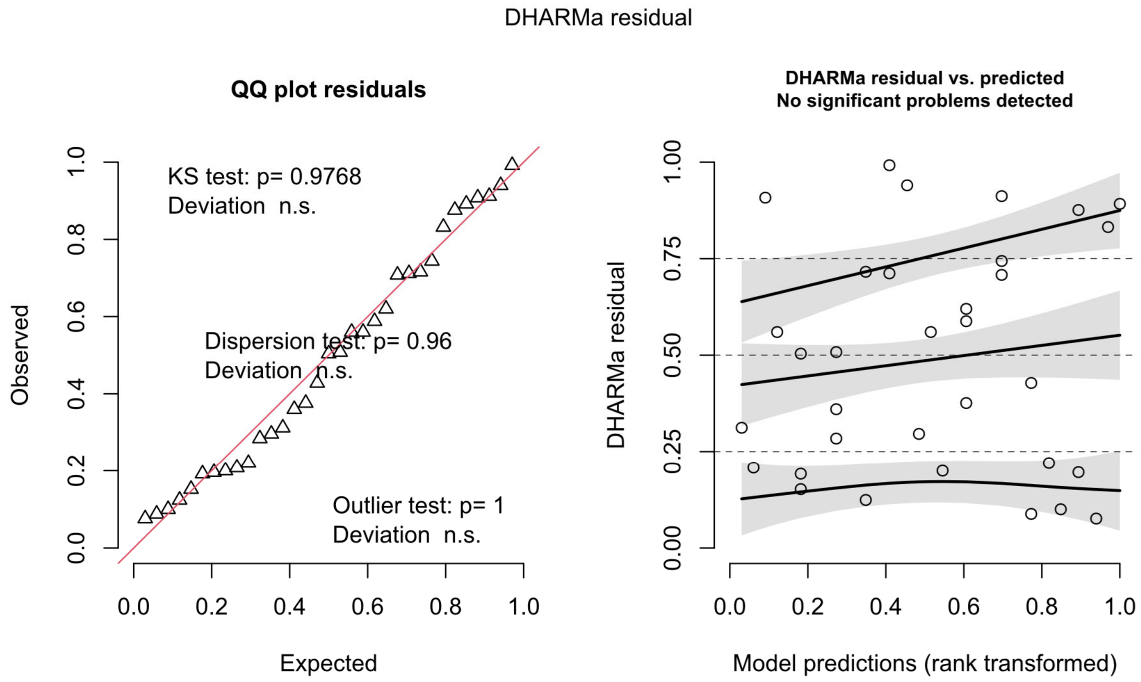


Figure S3. DHARMa simulated residuals for the final community size and age model, related to Table 2.
